# Space-based and object-based saccadic selection in visual working memory

**DOI:** 10.64898/2026.05.05.723053

**Authors:** Olga Shurygina, Laura Alexandra Wirth, Martin Rolfs, Sven Ohl

## Abstract

Saccades made during memory maintenance prioritize memory for the saccade target, but it is unclear if this benefit is specific to a location or extends across memorized objects. In three experiments, we examined whether saccadic selection spreads to other locations within the same object. In Experiment 1, we asked observers to remember three oriented Gabors presented either within contour-defined objects or without object structure. A subsequent movement cue prompted observers to move their eyes to the indicated location. We then probed memory for stimuli at locations equidistant from the saccade target, in either the same or a different object. Memory was best for stimuli at locations congruent with the saccade target, and consistently weaker for other stimuli presented in the same or a different object than the saccade target. In Experiment 2, we created more complex objects by adding more object features to the stimulus. Again, memory performance was best for stimuli congruent with the saccade target location, whereas memory in incongruent trials was worse and similar for stimuli in the same object as the saccade target and in a different object. In Experiment 3, we tested if saccadic selection is present and propagates within the object in a change detection task. Again, memory performance (i.e., change detection) was best at the saccade target location. However, this memory benefit also spread to other locations within the same object when the saccade target contained non-oriented noise. Our results imply that saccadic selection in visual working memory is primarily space-based but can also spread towards locations within the object where a saccade was directed.

## Introduction

Successfully navigating in situations such as a crowded airport terminal requires complex goal-directed movements based on visual information that can be retained and accessed even after the original sensory input is no longer available. For example, one might need to check whether a bag is still beside them after briefly shifting attention to a flight-information display. We maintain such visual representations in a capacity-limited visual working memory (Bays & Husain, 2008; Cowan, 2001; Luck & Vogel, 1997; Nakayama, 1991; Pashler, 1988; for reviews see Luck & Vogel, 2013; Ma et al., 2014). Because of this limited capacity, cognitive systems must select which information is encoded and preserved. Attention can be directed to space, features, objects, and time to select information for encoding into visual working memory or to prioritize information during memory maintenance (Griffin & Nobre, 2003; Kalogeropoulou et al., 2017; Landman et al., 2003; Wei et al., 2024; for review see Souza & Oberauer, 2016). Recent theories of visual working memory have emphasized the role of action in guiding this prioritization. Movements themselves—and even their preparation—can shape not only which parts of the visual scene are prioritized (Deubel & Schneider, 1996; Kowler et al., 1995; Li et al., 2016; Ohl et al., 2017; Rolfs & Carrasco, 2012; see Li et al., 2021 for review) but also which memory representations are maintained (Heuer, Ohl, & Rolfs, 2020; Myers et al., 2017; Olivers & Roelfsema, 2020; Van der Stigchel & Hollingworth, 2018; van Ede, 2020; van Ede & Nobre, 2023). Simple actions such as saccadic eye movements (Hanning et al., 2016; Hanning & Deubel, 2018; Ohl & Rolfs, 2017, 2018, 2020; Ohl, Kroell, & Rolfs, 2024), and reaching movements (Heuer et al., 2017; Heuer & Schubö, 2017), as well as their mere preparation (Trentin et al., 2023, 2024) prioritize information in visual working memory that aligns with current action goals. We will refer to the selection of memory contents through saccadic eye movements as *saccadic selection* in what follows.

Memory performance is best at the saccade target location even if this location is unlikely to be probed in a subsequent memory test (Ohl & Rolfs, 2017, 2020). In other words, saccadic selection in visual working memory occurs automatically. For visual perception it is well established that such automatic presaccadic attention spreads to other locations within the same perceptual groups (Wannig, Stanisor, & Roelfsema, 2011; Ghahghaei & Verghese, 2017; Shurygina, Pooresmaeili, & Rolfs, 2021) and within the same objects (Malcolm & Shomstein, 2015). Moreover, during prolonged fixation, attention guided by visual cues selects objects containing the cued location (Duncan, 1984; Egly et al., 1994; Jeurissen et al., 2016; for reviews see Scholl, 2001; Cavanagh et al., 2023). Similarly, prioritization in visual working memory guided by retro-cues extends to other locations within a cued object (Lin et al., 2021; Peters et al., 2015). However, it remains unknown whether saccadic selection in visual working memory also spreads beyond the saccade target location. Answering this question will allow us to determine the level at which saccades may select representations in visual working memory. If saccadic selection is purely space-based, the memory advantage would not spread beyond the saccade target location. In contrast, if saccadic selection operates at the level of maintained objects, the memory advantage would extend beyond the saccade target location, enhancing memory for all items within the object containing the saccade target.

Here, we tested whether saccades towards memorized items also improve memory for other items within the same perceptual object. In three experiments, we show that adding perceptual objects does not break saccadic spatial selection in either an orientation memorization task (**Experiments 1** and **2**) or a change detection task (**Experiment 3**). Furthermore, we demonstrate that selection spreads within an object given certain conditions in **Experiment 3**.

## Experiment 1

In **Experiment 1**, we asked participants to memorize three oriented gratings and make a cued saccadic eye movement after their disappearance (**Figure 1**). At the end of each trial, participants reported the orientation of one remembered stimulus. Importantly, the movement cue that prompted the saccade during memory maintenance was uninformative for the memory test. This experimental design was similar to the one used in previous studies to characterize saccadic selection in visual working memory (Ohl & Rolfs, 2017, 2018, 2020). In those studies, memory performance was better for stimuli presented at the saccade target and dropped at the neighboring locations, supporting the hypothesis that saccadic selection in memory is spatially specific (Ohl et al., 2024). In this experiment, we examined whether spatial specificity of saccadic selection in memory is modulated by object structure. We added an “object present” condition with outlines around the placeholders organizing them in two simple C-shaped objects (using the Gestalt law of common region; Palmer, 1992), keeping the rest of the experimental setup identical to the previous study design (**Figure 1**). We tested whether the memory benefit at the saccade target could spread to neighboring locations within the same object or whether the memory advantage would remain restricted to the saccade target location. The spread of performance benefits to other locations within the same object is a reliable way to quantify object-based attention (see reviews by Scholl, 2001, and Cavanagh et al., 2023). By contrasting the spatial specificity of saccadic selection in conditions with and without contour-defined objects, we can determine how the presence of objects alters saccadic selection in memory within our established paradigm.

**Fig. 1.**
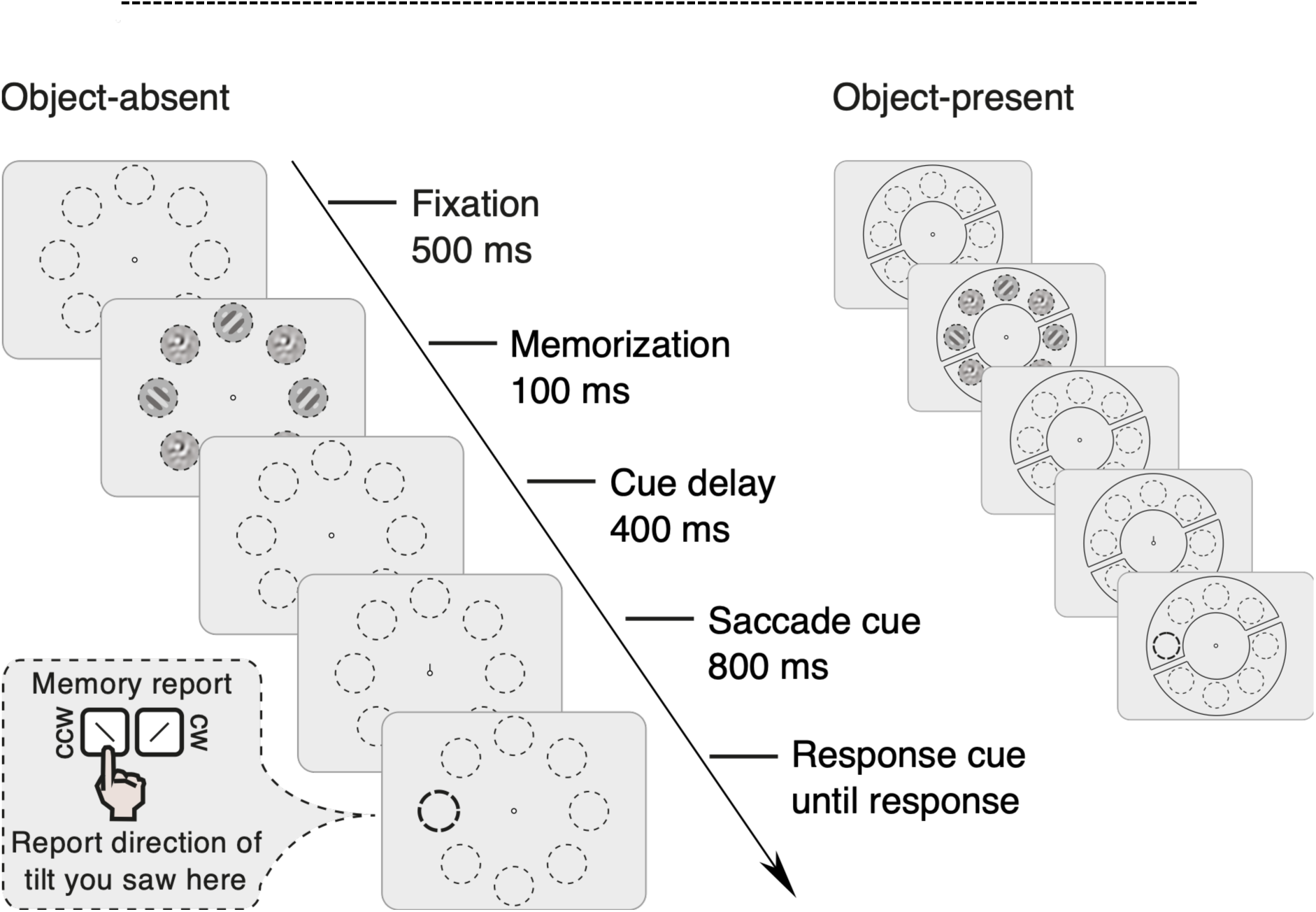
Experimental procedure of Experiment 1. We presented a memory array for 100 ms that contained three oriented Gabors and five noise patches, placed within placeholders (dashed outlines) on an imaginary circle. After a delay of 400 ms following array offset, a movement cue appeared at the fixation point, prompting observers to move their eyes to the indicated location. Another 800 ms after movement-cue onset, a probe highlighted one of the locations that previously contained an oriented Gabor. Observers reported the orientation (clockwise or counterclockwise from vertical) they remembered seeing at that location. In each trial, the saccade target and memory probe could be congruent (same location) or incongruent (different locations). The saccade target was uninformative about which Gabor would later be probed. In half of the trials, two black contours each surrounded four adjacent locations in the array, grouping them into two objects (object-present trials). In these trials, the probed location could be either in the same object or a different object than the saccade target. We randomly interleaved these trials with trials without contours (object-absent trials). In the exemplary trials above, the saccade target and the memory probe are incongruent.

### Methods

#### Participants

We tested 10 naïve participants (3 males, 7 females; ages 23–30, *M* = 26.3, *SD* = 2.26; all right-handed; 9 right-eye dominant, 1 left-eye dominant) in three 1-hour sessions with at least one night between consecutive sessions. The first session served as a training session to verify rapid saccade generation in response to a movement cue. No participant was excluded from the study. All participants had normal or corrected-to-normal vision and eye dominance was assessed using a hole-in-a-card test. Informed written consent was obtained prior to the experiment. Participants were compensated with 8.50€ per session plus a 2€ bonus upon completing all sessions.

#### Materials and procedure

Participants sat in a dark room at a viewing distance of 57 cm with their heads placed in a chin and forehead rest to reduce head movements. We recorded the positions of the observers’ dominant eye with an Eyelink 1000 Desktop Mount (SR Research, Ottawa, ON, Canada). For stimulus presentation we used a gamma-corrected VIEWPixx/3D (VPixx Technologies Inc., Saint Bruno, QC, Canada) in scanning backlight mode (luminance 0 to 100 cd/m^2^; pixel response time ∼1 ms) at a spatial resolution of 1920 × 1080 pixels and 120 Hz refresh rate. The experiment ran on a DELL Precision T3600 (Debian GNU Linux 8) and was implemented in MATLAB (The Math Works, Inc., 2020) using the Psychophysics Toolbox 3 (Brainard, 1997; Pelli, 1997) and the Eyelink Toolbox (Cornelissen et al., 2002).

Eye and screen coordinates were aligned using a standard nine-point calibration and validation routine at the beginning of each session, after breaks, and whenever recalibration was required. Before every trial onset, participants needed to direct their gaze at a fixation point (a white disk with a diameter of 0.17 degrees of visual angle (dva) enclosed by a black contour with a diameter of 0.68 dva and a width of 0.085 dva) directly at the center of the screen. A trial started as soon as the participants’ eye positions remained within a 1.5-dva circular region centered on fixation for at least 300 ms.

We presented stimuli on a gray background (luminance 77 cd/m^2^). The fixation point that we used for the fixation control at the beginning of the trial remained on the screen and in addition eight circular placeholders (black circle outline; 1.95 dva diameter) forming an imaginary circle at an eccentricity of 6 dva from the screen center appeared. The placeholders indicated the potential locations where the stimuli of the memory set could be presented. In half of the trials (object-present trials), the eight placeholders were grouped into two objects by displaying two additional contours (3 pixels wide; each consisting of two semicircles with inner borders 4.25 dva and outer borders 7.75 dva from fixation and two black lines connecting the semicircles, resulting in a closed object; contours enclosed 4 placeholders each). Placeholders and contours remained on the screen for the remainder of the trial.

After 500 ms, a memory array consisting of 3 oriented Gabor patches (+45° or -45° from vertical, 50% contrast, frequency of 2 cycles per degree) and five noise patches (pixel noise, band-pass filtered from one-half to twice the Gabors’ spatial frequency, 50% contrast, 0.65° SD Gaussian envelope) appeared for 100 ms inside randomly assigned placeholders. Observers memorized the Gabor orientations for a later test. Following a 400-ms delay, a movement cue appeared at fixation (0.26 dva black line segment extending beyond the fixation outline) prompting a saccade within 400 ms to the indicated placeholder. After an additional 800 ms delay, the response cue (i.e., thickening of one placeholder’s outline from 0.05 dva to 0.085 dva) instructed observers to report the remembered orientation (clockwise or counterclockwise from vertical) using keys N and B.

Importantly, in **Experiment 1** the movement cue always pointed to a placeholder that had contained a Gabor, but the cue was uninformative about which item would be tested. Thus, the saccade target location matched the memory-probe location in 1/3 of trials; in the remaining 2/3, a non-target location was probed.

We aborted trials online if a blink occurred, the gaze deviated more than 1.5 dva from fixation before movement-cue onset, or if the saccade latency exceeded 400 ms. Aborted trials were repeated in a randomized order at the end of each block.

Each block contained all conditions (saccade target and memory probe congruent vs. incongruent; objects present vs. absent; same vs. a different object trials). A test session consisted of 12 blocks of 48 trials, resulting in 1,152 trials across the two test sessions and 576 trials in the training session for each participant.

#### Data analysis

For saccade detection, we first transformed raw eye positions into 2D velocity space offline and classified successive eye-position samples as saccades if velocity exceeded the median by 5 SDs for at least 8 ms (Engbert & Mergenthaler, 2006). Saccadic events separated by less than 20 ms were merged. The first saccade landing within 3.6° of the saccade-target center (i.e., 60% of the target’s eccentricity) was classified as the response saccade. We excluded trials containing saccades with amplitudes larger than 1 dva prior to the response saccade, as well as trials with blinks, from further analyses. In total, 9,806 trials (85.1%) entered the final data analysis.

We conducted a repeated-measures ANOVA (rmANOVA) for statistical inference, followed by post-hoc paired t-tests for significant effects. Error bars are ±1 within-subject standard error of the mean (SEM), in the text we report 95% within-subject confidence intervals (CI; Baguley, 2011; Morey, 2008). For this and subsequent experiments, data and analysis scripts are available on the Open Science Framework (https://osf.io/2e7ay/overview?view_only=f4e78e29fb4e48c68ae9a68b0506654a).

### Results

In line with previous studies, memory performance was better at the saccade target as compared to other locations (Hanning & Deubel, 2018; Hanning et al., 2016; Ohl & Rolfs, 2017, 2018, 2020; Ohl et al., 2024; **Figure 2a**). The presence of objects (i.e., two black contours grouping adjacent locations into two objects) tended to decrease overall memory performance while the general memory benefit in congruent over incongruent trials was preserved. We corroborated this finding by a two-way (congruency: congruent vs. incongruent; object condition: present vs. absent) rmANOVA. We observed a significant main effect of congruency (*F*(1,9) = 28.89, *p* < 0.001) with better performance in congruent (proportion correct [pc] 0.84; 95% confidence interval [CI] [0.82, 0.86]) than in incongruent trials (pc 0.73; 95% CI [0.70, 0.76]). The presence of objects resulted in a marginally significant influence (*F*(1,9) = 5.08, *p* = 0.051)—showing a small decrease in memory performance in object-present trials (pc 0.75; 95% CI [0.74, 0.76]) as compared to object-absent trials (pc 0.78; 95% CI [0.77, 0.79]). There was no significant interaction between congruency and object presence (*F*(1,9) = 0.23, *p* > 0.250).

**Fig. 2.**
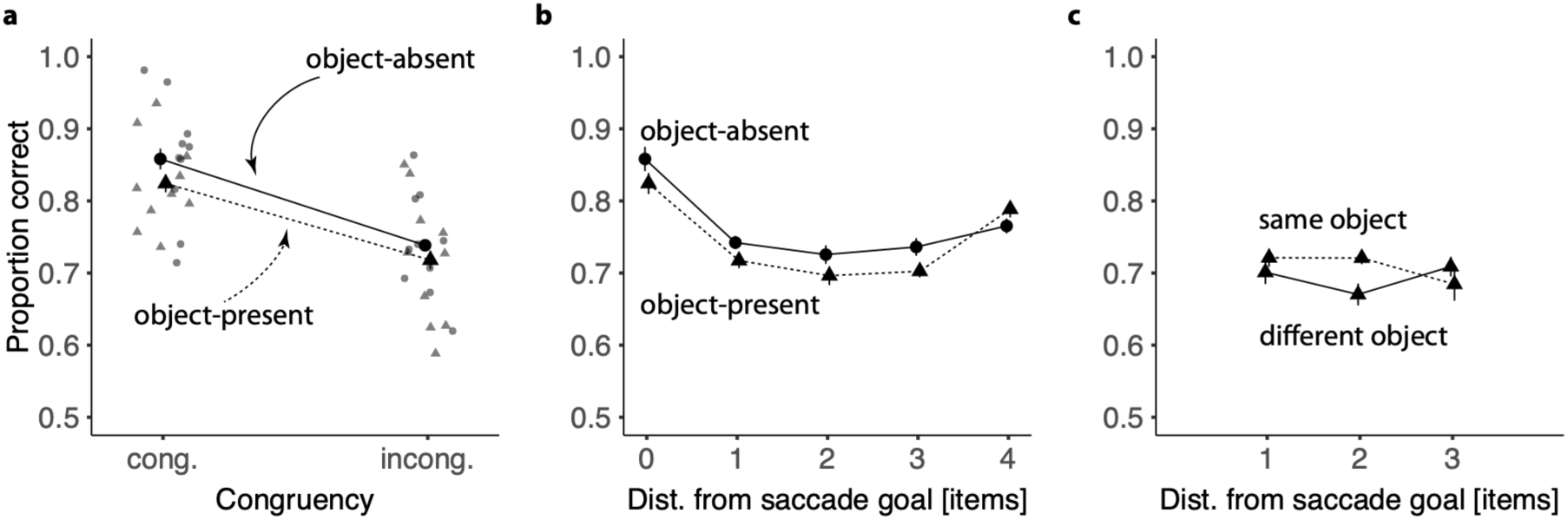
Results of Experiment 1. **a** Mean performance ±1 within-subject SEM as a function of congruency between the probe location and the saccade target (congruent vs. incongruent) in object-absent (circles, solid lines) and object-present trials (triangles, dashed). Corresponding gray symbols show individual participants’ performance. **b** Spatial specificity of saccadic selection in visual working memory. Mean performance ±1 SEM is displayed as a function of distance between the memory probe location and the saccade target location, separately for object-present and object-absent trials. **c** Assessing the same-object advantage by comparing memory performance ±1 SEM for locations that were either in the same or a different object than the saccade target location.

#### The spatial specificity of saccadic selection

We compared the spatial specificity of saccadic selection in object-present and object-absent trials (**Figure 2b**). In line with our previous findings, we observed best memory performance at the saccade target location and a sharp drop in memory performance already at the location next to it. Memory performance in object-present trials was consistently below performance in object-absent trials, apart from the most distant location. Here we found a small increase in memory performance as compared to the intermediate locations, and performance in object-present trials surpassed memory in object-absent trials. The finding was corroborated by a two-way (distances from saccade target: 0, 1, 2, 3, or 4 items; object condition: present vs. absent) rmANOVA. The effect of distance between the saccade target and the memory probe was highly significant (*F*(4,36) = 18.95, *p* < 0.001). Memory performance was better at the saccade target location than its neighboring location (Δpc_0-1_ 0.11; 95% CI [0.07, 0.16]). Memory performance at the incongruent locations was not statistically different between the intermediate locations (all *ps* > 0.150) but increased slightly at the location opposite the saccade target compared to its neighbor (*t*(9) = 5.47, *p* < 0.001; Δpc_4-3_ 0.06; 95% CI [0.034, 0.083]). Again, the main effect of objects in this rmANOVA showed only a trend (*F*(1,9) = 3.87, *p* = 0.081). The main effects were not accompanied by an interaction (*F*(4,36) = 1.32, *p* > 0.250), therefore showing similar spatial specificity for object-present and object-absent trials.

#### No same-object advantage for saccadic selection in memory

We assessed whether memory benefited from the presence of objects for stimuli that were presented in the same object as the saccade target location. To this end, we compared memory performance in object-present trials in which the saccade target and the memory probe were equidistant but located either within the same or in different objects. Numerically, memory performance for stimuli in the same object as the saccade target was slightly better than for stimuli in a different object at distances of 1 and 2 items, but not at 3 items (**Figure 2c)**. For inferential analysis, we ran a two-way (object location: same vs. different; distance from the saccade target: 1, 2, or 3 items) rmANOVA. However, the main effect of object location was not significant (*F*(1,9) = 0.58, *p* > 0.250). That is, memory performance was similar in incongruent trials irrespective of whether the probed stimuli were presented in the same object as the saccade target (0.72; 95% CI [0.694, 0.737]) or in a different object (0.70; 95% CI [0.676, 0.719]). The main effect for distance was also not significant for these intermediate locations (*F*(2,18) = 0.44, *p* > 0.250), nor was there an interaction (*F*(2,18) = 1.70, *p* = 0.211).

#### No speed-accuracy tradeoff and no difference in saccade characteristics

We did not observe a speed-accuracy tradeoff. A two-way (congruency: congruent vs. incongruent; object condition: object-present vs. object-absent) rmANOVA revealed a significant main effect of congruency (*F*(1,9) = 46.02, *p* < 0.001). Participants responded faster in congruent (501.9 ms; 95% CI [482.9, 520.63]) as compared to incongruent trials (743.1 ms; 95% CI [724.2, 762.0]). The main effect of object condition (*F*(1,9) = 0.34, *p* > 0.250) and the interaction were not significant (*F*(1,9) = 0.04, *p* > 0.250).

Finally, saccade latency and saccade amplitude did not vary across experimental conditions as revealed by two-way (congruency: congruent vs. incongruent; object condition: object-present vs. object-absent) rmANOVAs. Congruency had neither an influence on saccade latency (*F*(1,9) = 0.46, *p* > 0.250; congruent: 204.7 ms; 95% CI [203.7, 205.7]; incongruent: 205.3 ms; 95% CI [204.3, 206.3]) nor saccade amplitude (*F*(1,9) = 2.30, *p* = 0.164; congruent: 5.52 dva; 95% CI [5.50, 5.54]; incongruent: 5.50 dva; 95% CI [5.48, 5.52]). Likewise, there was no significant main effect of object presence on saccade latency (*F*(1,9) = 0.15, *p* > 0.250; present: 204.9 ms; 95% CI [204.1, 205.8]; absent: 205.3 ms; 95% CI [204.4, 206.1]), and no statistically significant influence on saccade amplitude (*F*(1,9) = 4.10, *p* = 0.074; present: 5.49 dva; 95% CI [5.46, 5.51]; absent: 5.53 dva; 95% CI [5.50, 5.55]). Also, we observed no significant interaction on saccade latency (*F*(1,9) = 0.001, *p* > 0.250) or saccade amplitude (*F*(1,9) = 2.63, *p* = 0.139).

### Discussion

In **Experiment 1**, we assessed whether saccadic selection in visual working memory was restricted to the saccade target location or whether memory benefits could also spread to memory representations that were part of the same object. In line with previous findings, we observed that saccadic selection was spatially specific to the saccade target location, and we did not find evidence for selection of other memory representations that were part of the same object. However, there was a numerical tendency towards higher memory performance in trials in which the saccade target and the memory probe were in the same object relative to trials in which they were equidistant but belonged to different objects.

There are multiple plausible reasons for this lack of a clear object-based advantage. One possibility is that saccades do not select entire objects in visual memory. Alternatively, aspects of our design may have reduced the likelihood of detecting such an effect. First, our object manipulation might have been too weak to make observers perceive the Gabor locations as strongly grouped or part of an object. Second, randomly interleaving object-present and object-absent trials could have allowed participants to suppress the contours and focus only on the oriented Gabors. We designed a second experiment to overcome these limitations.

## Experiment 2

In **Experiment 2**, we addressed the potential concerns raised in the previous experiment. First, we asked whether the presentation of objects in a blocked design would allow observers to more readily take advantage of processing the available objects and therefore reveal a clearer same-object advantage in memory performance. Second, we implemented additional Gestalt principles (Koffka, 1935, as reviewed in Wagemans et al., 2012; Köhler, 1929/1971; Metzger, 1936/2006; Wagemans, 2018; Wertheimer, 1938) that are known to strengthen the perception of objects, and compared whether more complex objects would produce a stronger same-object advantage than simple objects. While the simple objects (as in **Experiment 1)** were defined only by common region, the complex objects were additionally distinguished by outline color, texture, and onset location.

Again, we hypothesized that we would find higher memory performance in congruent trials (i.e., in trials in which the saccade target and the memory probe were at the same location) relative to incongruent trials (i.e., trials in which the saccade target and the memory probe were at different locations) in trials with simple and complex objects. Likewise, we hypothesized that memory performance would be higher for orientations presented within the same object as the saccade target compared to orientations presented at the same spatial distance from the saccade target but within a different object. We expected the same-object advantage to be more pronounced for complex objects than for simple objects.

### Methods

#### Participants

In total, 10 new, naïve participants (3 males, 5 females, 2 diverse; ages 18-27, *M* = 22.6, *SD* = 2.73; 9 right-handed, 1 left-handed; 9 right-eye dominant, 1 left-eye dominant) were tested in three 1-hour sessions with at least one night between consecutive sessions. The first session served as a training session, and the other two as test sessions. Two additional participants were excluded and replaced because their performance remained near chance level after the second session. Everyone was tested for normal or corrected to normal eyesight as well as for eye dominance using a hole-in-a-card test. Informed written consent was given prior to participation. Participants were compensated at 9€ per hour with a 3€ bonus for completing all three sessions.

#### Materials and procedure

The experimental procedure was largely identical to that of **Experiment 1** except for the following changes. First, the two trial types (here, simple vs. complex objects) were no longer interleaved but presented in blocks. We presented 6 blocks with simple objects and 6 blocks with complex objects, randomly interleaved in the test sessions.

Second, while the simple-object trials were identical to the object-present trials of **Experiment 1**, the complex-object trials included an additional 400 ms before memory array onset to allow for sequential object presentation. The first object appeared at its final position after 600 ms of fixation. The second object appeared simultaneously, moving from a corner of the screen to its final position along a straight-line trajectory over 300 ms. Participants maintained central fixation throughout.

Third, the object contours were no longer composed of two semicircles and two connecting lines but of 18 smaller line segments (3 replacing each original connecting line and 6 replacing each semicircle) forming an irregular polygon (**Figure 3**). The contour of each object was colored either blue or orange (RGB values with alpha: [0.1 0.3 0.7 1] and [0.65 0.30 0.15 1]), and all placeholders belonging to a given object were colored accordingly. Moreover, each object was filled with one of four texture categories (wood, washed and worn, vintage-photo-overlay, paper).

**Fig. 3.**
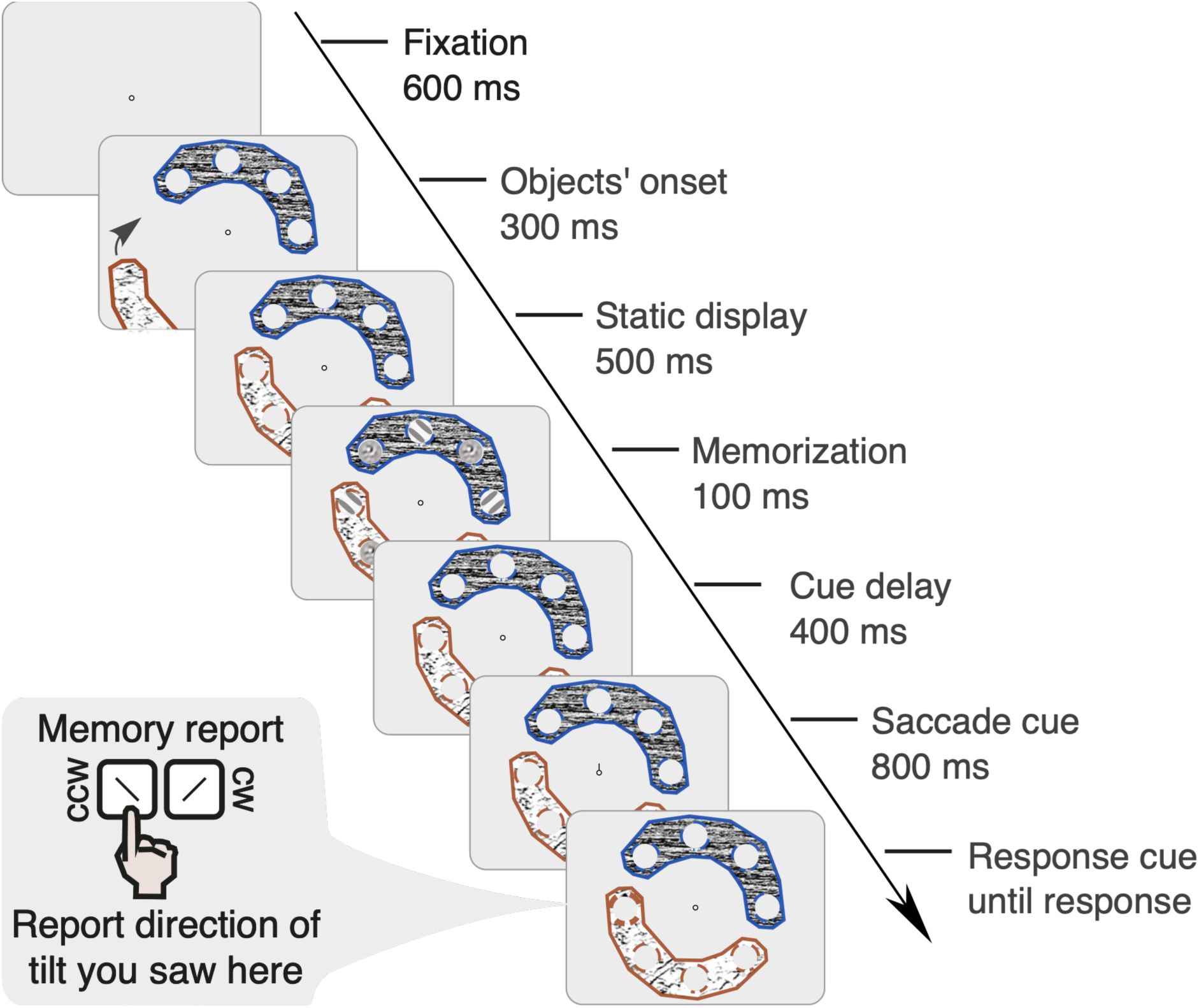
Experimental procedure of Experiment 2 (complex-object trial). We presented a memory array consisting of 3 oriented Gabors and 5 noise patches, placed on an imaginary circle, for 100 ms. 400 ms after the offset of the array, a movement cue appeared at fixation, instructing observers to saccade to the indicated location. Another 800 ms later, a probe highlighted one of the locations that previously contained an oriented Gabor, and observers reported the orientation they remembered seeing there (clockwise or counterclockwise from vertical). In each trial, the saccade target and memory probe could be congruent (same location) or incongruent (different locations). However, the saccade target was uninformative about which Gabor’s orientation would later be tested. In half of the blocks (shown here), we included additional Gestalt cues (i.e., color, texture, irregular borders, and separate onset and motion paths) to create complex objects. Trial duration was extended in the beginning prior to static display presentation. In all trials, the probed location could be part of the same or a different object.

#### Data analysis

For saccade detection we applied the same preprocessing steps as described in **Experiment 1**. In total, 10,189 trials (88.4%) entered the final data analysis of **Experiment 2.**

### Results

#### Similar saccadic selection for simple and complex objects

In **Experiment 2**, we again observed better performance in congruent trials compared to incongruent trials (**Figure 4a**)—an advantage of similar magnitude for both simple objects (Δpc 0.089; 95% CI [0.047, 0.130]) and complex objects (Δpc 0.095; 95% CI [0.062, 0.127]).

**Fig. 4.**
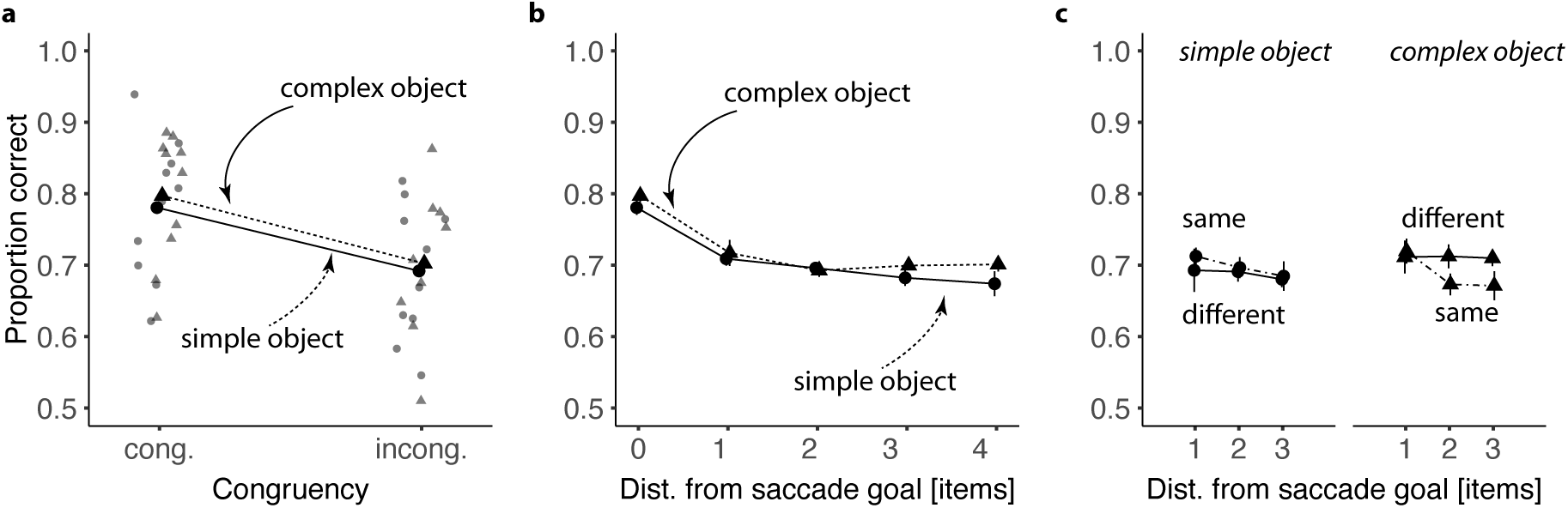
Results of Experiment 2. **a** Performance as a function of congruency (congruent vs. incongruent) in trials with simple (circles, solid lines) and complex objects (triangles, dashed). Corresponding gray symbols show individual participants’ performance. **b** Spatial specificity of saccadic selection in visual working memory. Performance is displayed as a function of distance between the memory probe location and the saccade target location, visualized separately for trials with simple and complex objects. **c** Performance for locations that were either in the same or a different object than the saccade target location, presented separately for trials with simple and complex objects. Means are shown with ±1 within-subject SEM.

For inferential analysis, we ran a two-way (congruency: congruent vs. incongruent; object type: simple vs. complex) rmANOVA. In line with our hypothesis, we found a significant main effect of congruency (*F*(1,9) = 55.02, *p* < 0.001), with better performance in congruent (0.79; 95% CI [0.78, 0.80]) than in incongruent trials (0.70; 95% CI [0.68, 0.71]). The main effect of object type was not significant (*F*(1,9) = 2.75, *p* = 0.132; complex objects: 0.73; 95% CI [0.72, 0.74]; simple objects: 0.72; 95% CI [0.71, 0.73]), nor was the interaction between congruency and object type (*F*(1,9) = 0.79, *p* > 0.250).

#### Similar spatial specificity for simple and complex objects

Next, we assessed the spatial specificity of saccadic selection for simple and complex object trials (**Figure 4b**). A two-way (distances: 0, 1, 2, 3 or 4 items away from the saccade target; object type: simple vs. complex) rmANOVA revealed a significant main effect of distance (*F*(4,36) = 17.91, *p* < 0.001), showing spatially specific saccadic selection. Performance was highest at the saccade target location and dropped sharply at the neighboring location (Δpc_0-1_ 0.077; 95% CI [0.042, 0.111]). Performance at intermediate incongruent locations did not differ (all *ps* > 0.160). Neither object type (*F*(1,9) = 1.59, *p* = 0.240) nor the interaction between distance and object type (*F*(4,36) = 0.26, *p* > 0.250) significantly influenced memory performance.

#### No same-object advantage irrespective of object complexity

We hypothesized that the more salient Gestalt cues in complex objects would yield a stronger same-object advantage compared to simple objects. However, memory performance did not differ between stimuli in the same vs different object at equal spatial distance from the saccade target, irrespective of object complexity (**Figure 4c**). A three-way (object location: same vs. different; distance: 1, 2, or 3 items away from the saccade target; object type: simple vs. complex) rmANOVA confirmed no main effect of object location (*F*(1,9) = 0.22, *p* > 0.250), no main effect for distance (*F*(2,18) = 1.94, *p* = 0.173), and no main effect of object type (*F*(1,9) = 0.11, *p* > 0.250). None of the interactions were significant (object location x distance: *F*(2,18) = 0.48, *p* > 0.250; object location x object type: *F*(1,9) = 2.27, *p* = 0.167; distance x object type: *F*(2,18) = 0.10, *p* > 0.250; object location x distance x object type: *F*(2,18) = 0.14, *p* > 0.250).

#### No speed-accuracy tradeoff and largely comparable saccade characteristics

As in **Experiment 1**, there was no evidence for a speed-accuracy tradeoff. A two-way (congruency: congruent vs. incongruent; object type: simple vs. complex object) rmANOVA revealed a significant main effect of congruency (*F*(1,9) = 74.01, *p* < 0.001), with faster responses in congruent (619 ms; 95% CI [586, 652]) than incongruent trials (871 ms; 95% CI [838, 904]). Object type did not influence response times (*F*(1,9) = 1.10, *p* > 0.250), nor was there a significant interaction (*F*(1,9) = 1.45, *p* > 0.250).

Similarly, saccade latency and saccade amplitude were largely comparable across conditions, as assessed with two-way (congruency: congruent vs. incongruent; object type: simple vs. complex object) rmANOVAs. Congruency had no effect on saccade latency (*F*(1,9) = 0.41, *p* > 0.250; congruent: 214.9 ms; 95% CI [213.8, 216.0]; incongruent: 214.3 ms; 95% CI [213.2, 215.4]) or amplitude (*F*(1,9) = 0.51, *p* > 0.250; congruent: 5.68 dva; 95% CI [5.66, .70]; incongruent: 5.68 dva; 95% CI [5.66, 5.70]). Object type had no statistically significant main effect on saccade latency (*F*(1,9) = 4.38, *p* = 0.066; simple: 216.8 ms; 95% CI [214.0, 219.5]; complex: 212.1 ms; 95% CI [209.4, 214.9]) or amplitude (*F*(1,9) = 0.02, *p* > 0.250; simple: 5.68 dva; 95% CI [5.59, 5.76]; complex: 5.69 dva; 95% CI [5.61, 5.77]) either. Finally, interactions were not significant for saccade latency (*F*(1,9) = 0.04, *p* > 0.250) but were significant for amplitude (*F*(1,9) = 7.70, *p* = 0.022). Overall, mean saccade amplitudes ranged from 5.64–5.72 dva, indicating very small absolute differences.

### Discussion

**Experiment 2** provides further support for the results from **Experiment 1**. Saccades again prioritized congruent representations in visual working memory in a highly spatially specific manner, and advantages did not spread to other representations within the same object (i.e., no same-object advantage). In **Experiment 2**, we used the same dual-task paradigm as in **Experiment 1** but included both simple and complex objects. Once more, we replicated the effect of spatial selection but found no evidence for object-based selection. Numerically, performance was slightly higher within the same object for simple objects, whereas the pattern was reversed for complex objects; these differences were not statistically reliable.

We re-examined the experimental design of **Experiments 1 and 2** to identify factors that may have favored space-based over object-based selection. First, the distinctive placeholders—visible throughout each trial—likely reduced the functional role of the objects, particularly because the memory probe always appeared within a placeholder. Second, the required saccades may have further increased attention to placeholders, as they were always directed toward one of them. Third, the Gabors used in the memory task were visually salient relative to the objects containing them—whether those objects lacked texture (simple objects in **Experiments 1** and **2**) or had a distinct texture (complex objects in **Experiment 2**). Consequently, both top-down factors (knowledge that both the saccade target and the probe always appeared within placeholders), and bottom-up factors (saliency differences between Gabors and objects) may have worked against object-based selection, thereby reinforcing spatially specific saccadic selection. To address these issues, we refined the experimental design and conducted **Experiment 3**.

## Experiment 3

In **Experiment 3**, we implemented several changes in the experimental design to address the factors that might have facilitated space-based saccadic selection in **Experiments 1** and **2**. We reduced the visual saliency of the memory array by embedding the memory array into the object texture. First, we removed all placeholders from the display. Second, we replaced the oriented Gabors with oriented texture patches derived from the object textures (1/f pink noise; see **Methods**). Third, to eliminate the saliency related to the saccade task, we balanced trials such that, in 50% of the trials, the movement cue pointed to a location within the object where no oriented item had been presented, while items were located elsewhere within the same object. We instructed participants to saccade in the direction of the movement cue toward the corresponding object region. Finally, to measure memory performance, we used a change detection task in which participants reported whether any item changed orientation after a retention interval (in 50% of trials, one item’s orientation changed within an object).

We hypothesized that, if object-based selection occurred, change detection would be more effective in trials where the saccade was directed to the region of the object containing the changed item. Additionally, we tested whether the space-based advantage would persist in this redesigned task, predicting the highest performance in trials in which the target and the changed item were congruent.

### Methods

#### Participants

In total, 10 new, naïve participants (3 males, 7 females; ages 18–38, *M* = 29.2, *SD* = 7.16; 9 right-handed, 1 ambidextrous; 8 right-eye dominant, 2 left-eye dominant) were tested in five sessions (75 min each) with at least one night between consecutive sessions. The first session served as a training session and the other four as test sessions. Fifteen participants were excluded after the training session because their performance was not substantially above chance or they did not make accurate saccades on time (we will discuss the reasons and implications of this unusually high drop-out rate below). Everyone was tested for normal or corrected-to-normal eyesight as well as eye dominance using a hole-in-a-card test. Informed written consent was given prior to participation. Participants were compensated with 10€ per hour.

#### Materials and procedure

The experimental equipment was identical to that used in **Experiments 1 and 2.** We aimed to keep the stimulus settings (brightness, eccentricities, colors) as similar as possible to those in **Experiment 2** (complex objects). The two objects appeared sequentially and moved to the central position of the screen. The shape, size and outline colors of the objects were identical to the complex objects in **Experiment 2**. Instead of using cutouts from different texture categories, however, we used pink-noise textures for the objects. This texture was pre-generated for each object at the beginning of each session, ensuring that the parameters remained constant across trials while avoiding repetition of identical textures. The most substantial change was the removal of the placeholders and the Gabors; instead, each object contained two orientation-filtered pink-noise patches derived from the object texture (see procedure in Kroell & Rolfs, 2022).

Participants were asked to maintain the two objects in memory and make a saccade in response to a movement cue during the memory-maintenance period (**Figure 5**). After a delay following the saccade, the objects reappeared on the screen, either with or without a subtle change in appearance (in 50% of the trials, one orientation-filtered patch changed orientation from clockwise to counterclockwise or vice versa). Participants reported whether they detected a change. As in **Experiments 1 and 2**, the movement cue prompting the saccade was uninformative for the memory task. We used the same online trial-termination criteria as in the previous experiments: trials were aborted if a blink occurred, gaze deviated more than 1.5 dva from fixation before movement-cue onset, or saccade latency exceeded 400 ms. Aborted trials were repeated in a randomized order at the end of each block. Each test session consisted of 15 blocks, 32 trials in each, resulting in 1,920 trials per participant. *Data analysis.* For saccade detection we applied the same preprocessing steps as in **Experiments 1 and 2**. In total, 13,582 trials (70.7%) entered the final data analysis of **Experiment 3.**

**Fig. 5.**
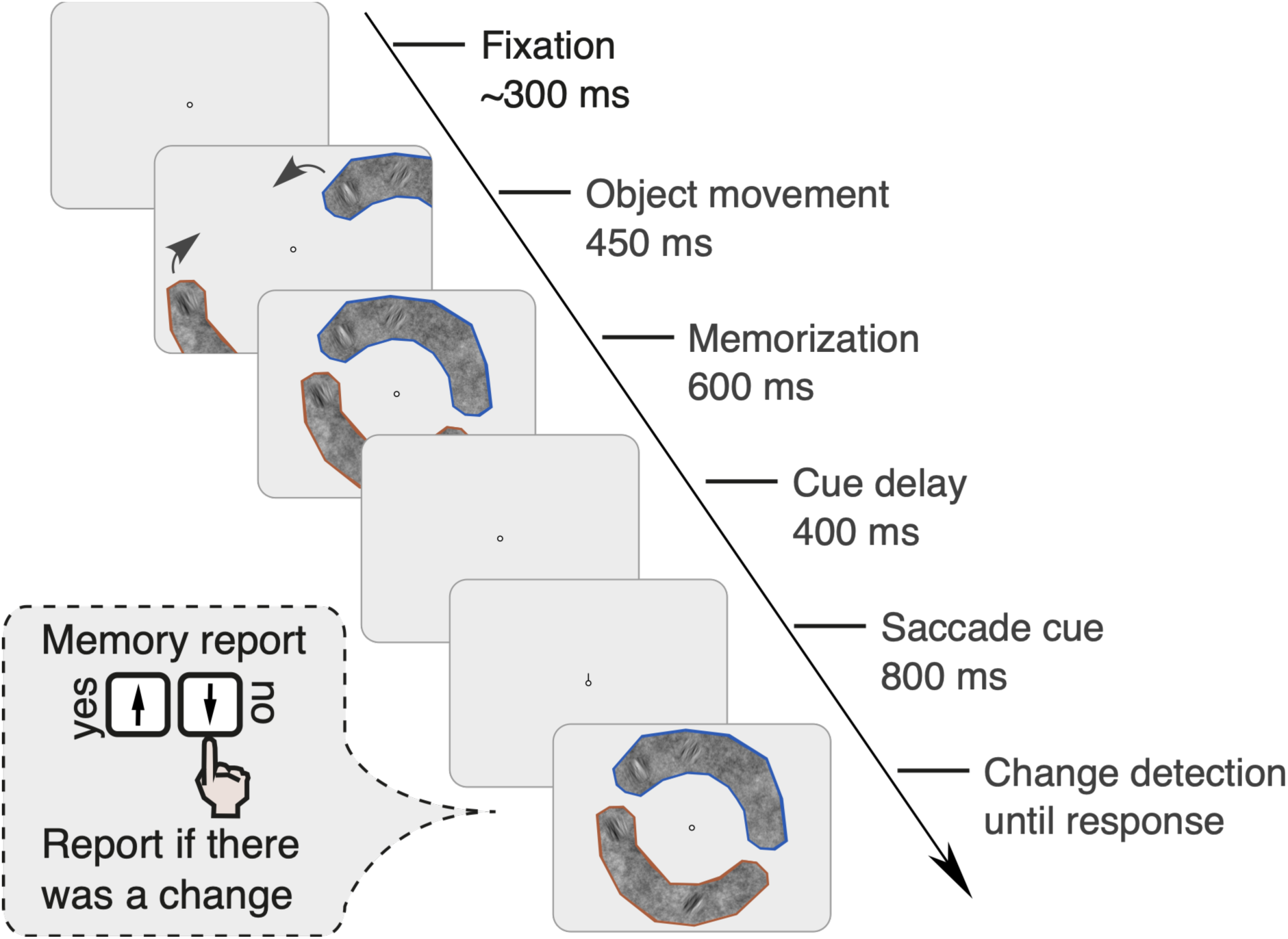
Experimental procedure of Experiment 3. The two objects appeared sequentially, moving from peripheral starting positions to their final locations along approximately straight trajectories. The second object began moving 150 ms after the first, with each movement lasting 300 ms. Both objects subsequently remained stationary for 600 ms before disappearing. The order of object presentation was randomized across trials. Following a delay of 400 ms after the offset of the array, a movement cue appeared at fixation, instructing participants to saccade to the indicated location. Another 800 ms later, a second stimulus was presented. In 50% of the trials, it was identical to the first stimulus (no-change trials) and in the other 50% of the trials, one of the orientation-filtered locations changed orientation (change trials). Participants reported whether they perceived a change.

### Results

Observers’ average accuracy in the task was 72.92% (95% CI [69.01%, 76.82%]). They responded “*no change*” on 60.60% (95% CI [55.54%, 65.67%]) of trials, which—despite differences in paradigm—is broadly consistent with classic findings on change blindness (Simons & Levin, 1997; Rensink et al., 1997). Overall sensitivity was d’ = 1.36 (95% CI [1.09, 1.62]). The mean proportions of hits, misses, correct rejections, and false alarms were 0.63 (95% CI [0.55, 0.70]), 0.38 (95% CI [0.30, 0.45]), 0.83 (95% CI [0.78, 0.89]), and 0.17 (95% CI [0.11, 0.22]).

For subsequent analyses, we focused on change trials only (50% of all trials), in which one item in the memory array changed orientation. This allowed us to compare performance based on the changed item’s (henceforth, the probe’s) spatial relation to the saccade target. We were specifically interested in whether performance differed when the changed item belonged to the same object as the saccade target versus a different object. We report the proportion of correct responses rather than d′, which is commonly used in change detection tasks. In our design, however, false alarm rates could not be calculated separately for each condition. The no-change trials (50%) could not be divided into the same categories as the change trials (e.g., congruent vs. incongruent, same vs. different object, distance from the saccade target), because these saccade-related distinctions are not meaningful when the display remains unchanged (e.g., it is not meaningful to determine whether no-change occurred in the same object as the saccade target location). As a result, condition-specific false alarm rates could not be estimated, and we compared proportions of correct responses directly.

Notably, unlike in the previous experiments, the movement cue appeared on a blank screen, and the cued location could have contained either a pink-noise field or an oriented patch, both embedded within the object. This enabled us to test whether saccades directed toward noise vs. oriented content differentially influenced memory performance, the same-object advantage, or saccade execution.

#### Saccadic selection in a change detection task

First, we assessed whether saccadic selection in visual working memory was preserved in the change detection task. Consistent with **Experiments 1** and **2**, change detection performance was significantly higher (*t*(9) = 5.96, *p* < 0.001; Δpc = 0.12; 95% CI [0.08, 0.17]) when the probe appeared at the saccade target (pc 0.73; 95% CI [0.71, 0.75]) than at all other locations (pc 0.61; 95% CI [0.59, 0.63]) (**Figure 6a**).

**Fig. 6.**
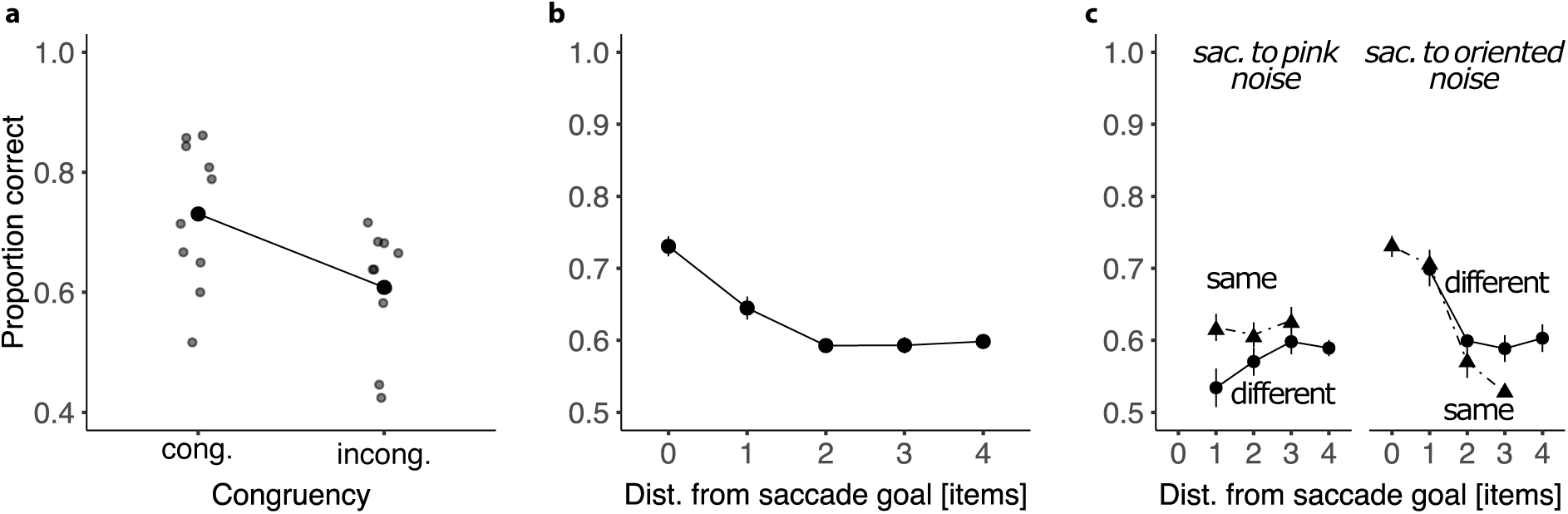
Results of Experiment 3. **a** Change detection performance in change trials as a function of congruency between the probe location and the saccade target (congruent vs. incongruent). Gray symbols show individual participants’ performance. **b** Spatial specificity of saccadic selection. Performance is shown as a function of the distance between the memory probe location and the saccade target location. **c** Same-object advantage. Change detection performance for probes appearing in the same versus a different object relative to the saccade target location. Data are shown separately for trials in which the saccade was directed toward pink noise versus orientation-filtered noise. Means are shown with ±1 within-subject SEM.

#### Spatial specificity in the change detection task

Change detection performance varied significantly with the distance between the changed item and the saccade target (*F*(4,36) = 14.88, *p* < 0.001). Performance peaked when the change occurred at the saccade-target location. Performance was significantly higher at this location than at the immediately adjacent one (*t*(9) = 3.23, *p* = 0.021; Δpc_0-1_ 0.09; 95% CI [0.03, 0.15] – p value is Holm-adjusted).

Unlike in our previous experiments, however, performance also improved at the location next to the saccade target, suggesting a more gradual spatial gradient of facilitation. Performance at that location was significantly higher than at the next location beyond the adjacent one (*t*(9) = 2.40, *p* = 0.040; Δpc_1-2_ = 0.05; 95% CI [0.00, 0.10] – p value is Holm-adjusted). Performance remained relatively stable at distances 2 to 4.

#### Same-object advantage

Next, we addressed our main question whether change detection performance in **Experiment 3** revealed a same-object advantage for the object containing the saccade target. We compared trials in which the changed item was embedded in the same object as the saccade target versus a different object. In the analysis, we statistically controlled for the distance between the saccade target and the location of the changed item, as well as for cue direction (saccades directed toward non-oriented pink noise vs. orientation-filtered noise patches).

We conducted a three-way rmANOVA with the factors object location (same vs. different), distance (1, 2, or 3 items), and cued location (non-oriented pink noise vs. oriented patch). The effect of distance was significant (*F*(2,18) = 4.37, *p* = 0.028), replicating the finding that performance at incongruent locations was highest at the location adjacent to the saccade target. There was no main effect of object location (*F*(1,9) = 0.344, *p* >0.5), and no main effect of cued location (*F*(1,9) = 2.35, *p* = 0.160). Crucially, we observed a significant interaction between object location and cued location (*F*(1,9) = 7.42, *p* = 0.023). A significant same-object advantage emerged when saccades were directed to a location previously occupied by non-oriented pink noise (Δpc = 0.05; 95% CI [0.01, 0.10], *t*(9.0) = 2.46, *p* = 0.036) (left panel in **Figure 6c**). No same-object advantage was observed when the saccade targeted a location that previously contained an orientation-filtered noise patch (Δpc = –0.03; 95% CI [–0.09, 0.03], *t*(9.0) = -1.01, *p* = 0.340).

#### Saccade parameters were affected by the visual content at the target location

Given that a same-object advantage emerged only when participants directed saccades toward non-oriented pink noise, we next tested whether saccade parameters were similarly influenced by the type of stimulus previously occupying the target location. Specifically, we examined whether saccade latency and amplitude differed between saccades directed to locations that previously contained an orientation-filtered noise patch versus non-oriented pink noise. As a sanity check, we verified that object location did not affect saccade metrics.

We conducted two separate two-way rmANOVAs testing the impact of cued location (non-oriented pink noise vs. orientation-filtered noise patch) and object location (same vs. different object for saccade target and probe) on saccade latency and amplitude. Cued location did not significantly affect saccade latency (*F*(1,9) = 3.23, *p* = 0.11). However, it did significantly influence saccade amplitude (*F*(1,9) = 9.82, *p* = 0.012). Saccades directed toward the former location of the pink noise were shorter than those directed to former locations of orientation-filtered patches (5.78 vs 6.06 dva; *M* = –0.28 [–0.48, –0.08], *t*(9.0) = –3.13, *p*=0.012). As expected based on previous experiments, object location had no effect on either parameter (all *p*s >0.1).

### Discussion

We showed that the type of visual content previously occupying the saccade target location systematically influenced both saccade amplitude and the same-object advantage in memory. Saccades were more accurate—reflected in larger amplitudes—when directed to locations that had previously contained orientation-filtered noise patches compared to locations previously occupied by non-oriented pink noise. Strikingly, however, an object-based advantage in change detection emerged only when saccades were directed toward the former locations of non-oriented pink noise. No such advantage occurred when the saccade targeted a former orientation-filtered patch. This dissociation suggests that the visual characteristics of the saccade target determine not only the oculomotor response but also whether object-based selection in visual working memory is engaged.

## General Discussion

In the present study, we contrasted two possible levels at which saccades may select representations in visual working memory. First, saccadic selection in visual working memory could be space-based, predicting that the memory advantage is confined strictly to the saccade target location. Second, saccadic selection could take place at the level of maintained objects, in which case the memory advantage would extend beyond the saccade target location, enhancing memory for all items within the object containing the saccade target. Our findings from **Experiments 1 and 2** support strong space-based saccadic selection in visual working memory, whereas the change detection task in **Experiment 3** revealed that saccadic selection in visual memory can also be object-based, in addition to robust space-based selection.

The finding from **Experiment 3** that saccadic selection in memory can in principle be object-based in addition to space-based is in line with other studies reporting object-based spread of attention in both perception and memory. For instance, reaction times are shorter when attending to locations in memory that were parts of attended objects, compared with equidistant locations in different objects. Moreover, brain activation in visual areas V1-V4 and parietal cortex is substantially higher for positions along the attended object, suggesting that attentional selection in working memory activates the entire object (Peters et al., 2015). Memory updates are faster for two equidistant colors on the same object than on different objects, and accuracy in retrieving memorized features is higher for un-cued features in the same object than for cued features in a different object (Lin et al., 2021).

Typically, such studies employ retro-cues in their experimental designs, which can be used strategically to suppress non-cued items or enhance cued memory representations (for review see Souza & Oberauer, 2016). In contrast, our experiments relied on saccades that were uninformative for the memory task, suggesting that, under certain conditions, saccadic selection can elicit an automatic spread of attention (**Experiment 3**). This aligns with findings in the perceptual domain showing that attention can spread automatically to locations grouped with the saccade target, even when those locations are irrelevant to the task. Such effects have been demonstrated both in neural activation patterns in primary visual cortex in monkeys (Wannig et al., 2011) and in increased visual sensitivity in humans (Shurygina et al., 2021).

### Factors modulating object-based selection

We next consider the factors that might have hindered or enabled object-based attention in our three experiments. In the first two experiments, in which participants had to recall whether an orientation was clockwise or counterclockwise, we observed a strong space-based effect: Memory performance improved at the saccade target location but did not extend to other parts of the object containing the target. In contrast, the third experiment, which used a change detection task, revealed both space-based and object-based advantages. Notably, the object-based effect emerged only when saccades were directed to regions containing a non-oriented texture, suggesting that both task demands and stimulus configuration critically influence object-based selection.

First, the change detection paradigm may have promoted a global encoding of the objects, encouraging participants to process entire objects rather than focusing narrowly on individual probes. In contrast, the delayed discrimination task in **Experiments 1 and 2** may have biased participants toward probe-specific encoding, allowing them to ignore object-level structure. Recently, assessing presaccadic attention in a change detection task revealed that visual sensitivity did not increase at the saccade target location (Gupta & Sridharan, 2024). Instead, the change detection task revealed a criterion shift leading participants to report changes at the saccade target location more often. In our study, we observed higher change detection rates at the saccade target location in a change detection task. However, because all locations shared the same false alarm rate, we could not separate sensitivity from a location-specific criterion. We avoided asking participants to localize changes, aiming to encourage holistic object perception, which may have resulted in overall better performance at the saccade target. Thus, our result is consistent with the general notion that the unit of selection in visual working memory can depend on task demands (Cao & Deouell, 2025).

Second, the stimulus features in **Experiment 3**—probes embedded within object textures, absence of placeholders, and irregular shapes with distinct outlines—likely enhanced perceptual grouping, making the objects more salient and cohesive while simultaneously reducing the saliency of individual probes.

Studying object-based selection inevitably raises the question: what constitutes an object in the observer’s perception? Gestalt principles define features that promote perceptual grouping (Van Geert et al., 2023). In previous work, strengthening perceptual grouping facilitated the spread of presaccadic attention to elements within the same group, even when those elements were task-irrelevant (Shurygina et al., 2021). In **Experiment 2**, we attempted to enhance perceptual grouping relative to **Experiment 1** by introducing distinctive textures, variable motion trajectories and onset locations, colored outlines, and irregular object shapes, yet memory performance remained confined to the saccade target location. We suggest that the placeholders caused probe elements to be perceived and encoded as separate items, such that object-level information could be disregarded as irrelevant. Placeholders can also induce masking or crowding effects, reducing perceptual capacity across the stimulus display (Puntiroli et al., 2018; Hanning & Deubel, 2022). In such cases, presaccadic attention shifts overcome the detrimental effect of crowding. Indeed, when placeholders were absent during saccade preparation, Puntiroli et al. (2018) observed no strong presaccadic attentional enhancement.

In **Experiment 3**, the absence of placeholders and embedding of probes within object textures allowed the objects to be perceived as cohesive units, while probes became less visually distinct. Under these conditions, we observed both a space-based enhancement of memory at the saccade target and a spread of this enhancement within the object containing the target. These results suggest that object-based selection in memory is conditional and task-dependent.

Interestingly, saccade parameters also differed depending on the visual content at the target. Saccades to non-oriented pink noise had shorter amplitudes and were less accurate than saccades to orientation-filtered probes. This suggests that remembering an oriented item at the saccade target location focuses selection, while saccades toward non-oriented regions allow memory selection to spread more broadly across the object. Analogous findings in perception show that performance benefits at a saccade target can extend across a textured region if it separates the saccade target from the background but rapidly decline with distance when the target is on a uniform background (Ghahghaei & Verghese, 2017).

### Robustness of space-based selection vs. fragility of object-based selection

Overall, our results suggest that object-based selection in memory is more fragile and task-dependent than the robust space-based selection, which consistently enhances memory at the saccade target. This aligns with a broader body of literature on object-based attention. First, space-based attention tends to be stronger and steadier than object-based attention (Egly et al., 1994; Francis & Thunell, 2022). Moreover, object-based effects are dependent on spatial direction and task type, often being stronger along the horizontal axis (Hein et al., 2016; Al-Janabi & Greenberg, 2016; Barnas & Greenberg, 2019; Chen & Cave, 2019) and in detection or comparison tasks rather than identification tasks (Pilz et al., 2012; Zheng & Moore, 2021). Spatial cue presence and validity also matter: Lower cue validity reduces the likelihood of observing an object-based effect (Lou et al., 2022), and according to Donovan et al. (2016), in the absence of a spatial (endogenous) cue, objects do not guide attention. Finally, while an increase in visual sensitivity is a gold standard for behavioral measures of attention, behavioral measures of object-based attention are often observed in reaction times rather than accuracy (Egly et al., 1994; Jeurissen et al., 2016). We suggest that similar principles may apply in visual working memory. As evidence, behavioral and electrophysiological studies show that object integration at early perceptual stages influences the processing at subsequent stages, facilitating the maintenance of individual objects and their constituent features, and leading to enhanced precision of representations stored in memory (Chen et al., 2024; Xie et al., 2021). Thus, unlike space-based selection, object-based selection in memory depends on an interplay of stimulus configuration, spatial orientation, and task demands.

### Limitations of the current study and future directions

We argued that the emergence of an object-based effect in **Experiment 3** was driven by changes to both the task and the stimulus. However, these changes introduced a new challenge: a high drop-out rate among participants. Prior to data collection, we defined exclusion criteria based on task performance and saccade accuracy. Specifically, participants who failed to reach above-chance accuracy in the first session were excluded from the experiment, as were participants who failed to execute accurate or timely saccades. In total, we excluded (and replaced) 15 participants in **Experiment 3**.

This contrasts sharply with the other two experiments: **Experiment 1** had no excluded participant, and **Experiment 2** excluded only two participants due to poor memory performance. The high exclusion rate in **Experiment 3** presents a notable limitation for the generalizability of our findings and reflects the increased difficulty of the task and stimulus setup. There are several potential ways to improve task accessibility without compromising the integrity of the design. One straightforward approach is to reduce the delay between the two stimuli, which should markedly increase change detection performance.

Importantly, several participants who failed to meet performance criteria reported that they could barely distinguish the oriented patches embedded in the noise texture. This suggests that the difficulty was not solely memory-related but may also stem from perceptual limitations. To address this, a promising follow-up design could incorporate an adaptive staircase procedure to individually adjust the contrast between oriented patches and the background texture. This approach can ensure that the items are perceptually discriminable for each participant (e.g., Kroell & Rolfs, 2022). However, this solution is non-trivial: the contrast must remain subtle enough that the oriented patches are not perceived as distinct objects, which could reintroduce item-based encoding strategies.

An alternative, and potentially simpler, solution would be to reduce the size of the stimulus display, thereby decreasing the distance between fixation and probe locations. This could enhance both perceptual encoding and saccadic accuracy, improving overall task feasibility. Another promising direction for future research is to move experimental designs toward more realistic, real-world scenarios because real, tangible objects elicit different perceptual, memory, and attentional responses than images (for review see Snow & Culham, 2021).

## Conclusion

Saccadic selection in visual working memory is robustly space-based, consistently prioritizing the saccade target location. Object-based selection, in contrast, can emerge under conditions that promote global object perception and reduce item saliency. These findings reveal that what we remember is shaped not just by where we look, but also by how our visual system organizes and prioritizes objects in memory.

